# Discovery and Repurposing of Pharmaceutical Agents as Antifungals: *In vivo* activity against *Coccidioides*

**DOI:** 10.1101/2024.02.21.581462

**Authors:** Heather L. Mead, Michael Valentine, Holly Yin, George R. Thompson, Paul Keim, David M. Engelthaler, Bridget M. Barker

**Affiliations:** The Translational Genomics Research Institute (TGen), Flagstaff, AZ, USA; Beckman Research Institute of the City of Hope; Department of Internal Medicine, Division of Infectious Diseases, and Department of Medical Microbiology and Immunology, University of California-Davis, Sacramento, California, USA; The Pathogen and Microbiome Institute, Northern Arizona University, Flagstaff, AZ, USA

**Keywords:** Coccidioidomycosis, Valley Fever, antifungal therapy, fungal infection, antifungal therapy, adjunctive agent, *Coccidioides*, sertraline

## Abstract

There is significant interest in identifying improved treatments for coccidioidomycosis, an endemic fungal infection found in the southwestern United States, and Central and South America. The current standard of care, fluconazole, often fails to completely eradicate *Coccidioides* infection; however, the cost of identifying new compounds is often high in terms of both finances and time. Therefore, repurposing existing pharmaceutical agents is an attractive option. In our previous work, we identified several compounds which inhibited fungal growth *in vitro*. Based on these findings, we screened a subset of these agents to increase the performance of fluconazole in a combination therapy approach, as compared to fluconazole alone, in a murine model. We observed that combination therapy of sertraline:fluconazole significantly reduced the amount of live fungus in the lung compared to fluconazole alone. These results suggest that sertraline can be repurposed as an adjunctive agent in the treatment of this important fungal disease.

## INTRODUCTION

*Coccidioides* spp. are pathogenic environmental fungi responsible for considerable morbidity and mortality within the United States^1^. These organisms cause the disease coccidioidomycosis (Valley Fever [VF]). Clinical manifestations vary depending upon both the extent of infection and the immune status of the host. Although many infections are of subclinical severity, some patients develop flu-like symptoms (*e.g*., cough, fever, chills, headaches, joint pain, rash) consistent with respiratory infection, while others develop extrathoracic dissemination requiring long courses of antifungal treatment. Patients with severe coccidioidomycosis are treated with an amphotericin B formulation, which carries significant potential toxicity and is currently available only intravenously. Those with mild to moderate infection are treated with triazole antifungals (fluconazole, itraconazole, voriconazole, posaconazole, or isavuconazole)^2^. These agents have exhibited efficacy against *Coccidioides* in both animal and human studies; however, these agents may have toxic effects, considerable drug-drug interactions exist, and refractory disease may persist despite treatment, mandating the continued search for additional therapeutic options^3^.

In contrast to traditional drug development pathways, the repurposing (or repositioning) of old drugs for new indications (such as antifungals) may significantly reduce the time, effort and costs required to bring new agents to clinical trials^4^. Recent work has shown that repurposing is a viable strategy, with synergistic *in vitro* activity leading to clinical trials evaluating the efficacy of combination therapy in cryptococcal infection. Previously, our team demonstrated the ability of dozens of compounds from the LOPAC small molecule library to inhibit *Coccidioides* growth *in vitro* (U.S. Patent 11,446,353). From these *in vitro* findings, we identified a subset of compounds which could be administered orally, were likely to have acceptable mammalian toxicity profiles, and may demonstrate adjunctive potential when combined with fluconazole, the recommended first line of treatment for VF. Here we evaluate the efficacy of these small molecules in a murine model of infection.

## Methods

### In vivo assays

Infection was established using 1,000 arthroconidia suspended in sterile PBS from *Coccidioides posadasii* strain Silveira (ATCC 28868) instilled intranasally under anesthesia with ketamine/xylazine (80 mg/kg/8mg/kg) injected intraperitoneally. Fungal cultures were grown, harvested and quantified as previously described^5^. CD1 male (n=10 per group) mice between the ages of 4-6 weeks were given tamoxifen (Sigma) in peanut oil, administered via oral gavage, for 3 days prior to fungal infection at 200 mg/kg/day^6^ or sertraline hydrochloride (Sigma) in saline, via oral gavage, for 6 days prior to fungal infection at 15 mg/kg/day. From 24 hours post-infection, animals in the treatment groups were dosed, via oral gavage every 24 hours, with: tamoxifen (200 mg/kg/day), sertraline (15 mg/kg/day), fluconazole (Terriquez) (15 mg/kg/day), vanoxerine (Sigma) (10 mg/kg/day); or combinations of tamoxifen:fluconazole (15:200 mg/kg/day), sertraline:fluconazole (15:15 mg/kg/day), or vanoxerine:fluconazole (10:15 mg/kg/day). Supplemental Table 1 outlines experimental groups. Mice were euthanized 6-8 days post infection, and lungs and spleens were harvested. Post-necropsy fungal burden was determined by homogenizing the entire lung or spleen, respectively, in 1 mL of sterile PBS. Serial dilutions of each homogenate were plated in duplicate on 2xGYE (2% glucose, 1% yeast agar) and incubated for 3-4 days at 30°C. Colony forming units (CFU) were counted and averaged between dilutions and replicates as described^5^. Plates were retained for a total of two weeks to observe for possible delayed growth.

### Statistical analysis

Statistical analysis and graphical representations were conducted using RStudio software R version 4.2.1^7^. A negative binomial regression model (MASS package^8^) was used to determine the reduction in log fungal burden.

## RESULTS

### Tamoxifen:fluconazole treatment decreases fungal burden and contributes to weight loss

The planned terminal date for each experiment was eight days post infection. However, all mice treated with either tamoxifen alone or in combination with fluconazole experienced a loss of >20% of total body weight, and were therefore euthanized on day 6. For the sake of statistical comparison, mice from the other groups (untreated infection, fluconazole treatment and control mice) were also euthanized on day 6 instead of day 8. Fungal burden in the lung was measured as the average CFU grown from whole lung homogenates in 1 mL of PBS. The untreated infection group had 1 × 10^6^ (SD 1.2 × 10^6^ cells/mL), fluconazole alone 7 × 10^3^ cells/mL, tamoxifen alone 5.3 × 10^5^ (SD 1.5 × 10^5^ cells/mL), and tamoxifen:fluconazole 4 × 10^3^ (SD 4.6 × 10^3^ cells/mL) total fungal colonies. Using a negative binomial model, combination therapy was observed to significantly reduce the log fungal burden in the lung compared to fluconazole alone (log reduction -2.46, *P* < 0.001, Table 1). We observed dissemination to the spleen in three groups: 50% untreated infected, 100% fluconazole, and 77% of tamoxifen-treated mice. No dissemination at all was observed in the combination therapy group. Uninfected mice gained weight (median, IQR; 1.55g, 0.75) over the duration of treatment, whereas untreated infected mice lost weight (median, IQR; -2.65g, 0.85). Tamoxifen:fluconazole treated mice lost weight during infection (median, IQR; -1.7g, 2.3) as described above; one fluconazole mouse was sacrificed on day 6 (gained 1g); and the remaining fluconazole + infection mice were sacrificed on day 8 (median, IQR; -0.6g, 1.05).

**Figure 1.**
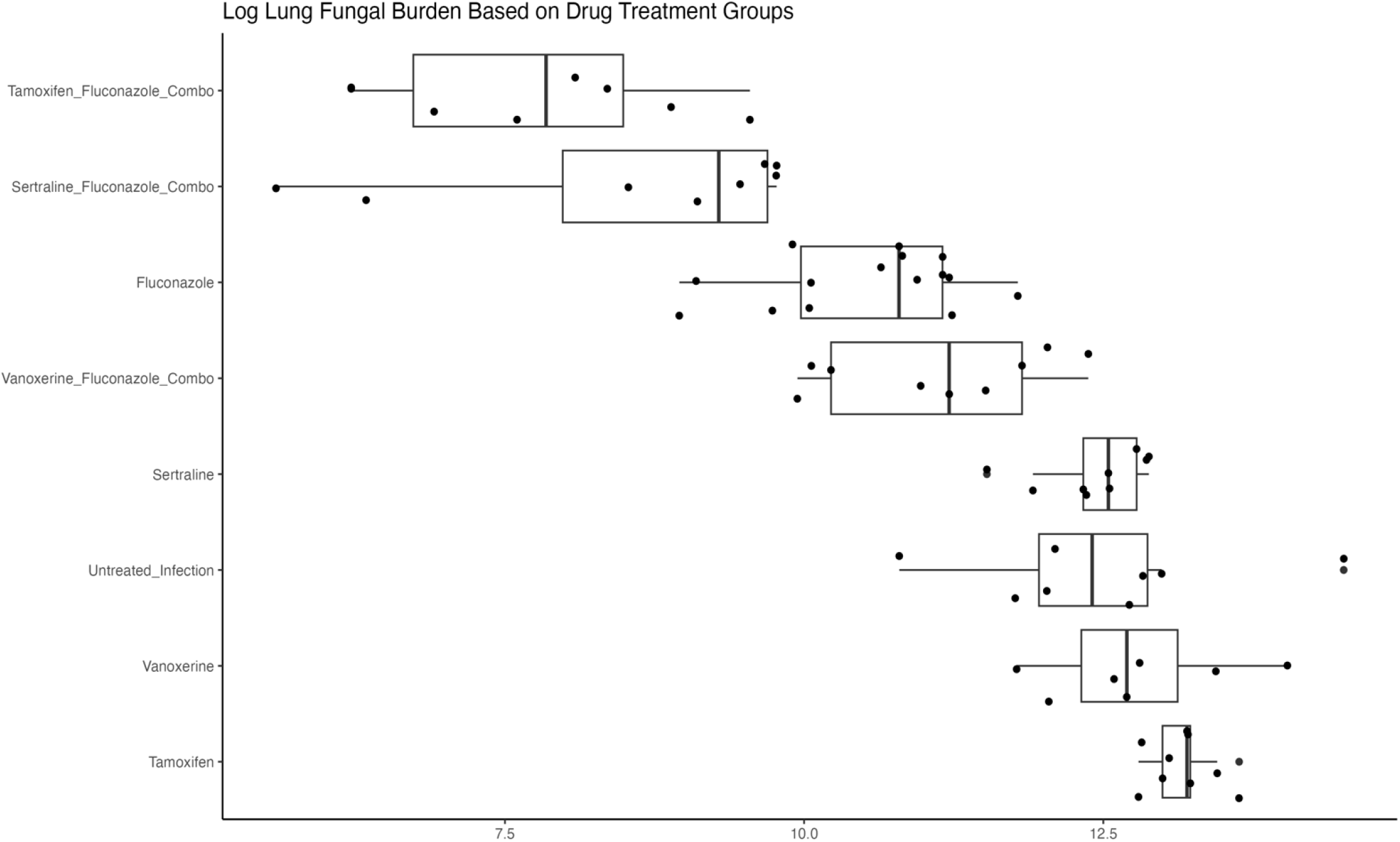
Combination tamoxifen:fluconazole and sertraline:fluconazole therapy significantly reduce fungal burden compared to fluconazole alone (p < 0.001). A negative binomial was used to compare the weighted average fungal burden in the lungs for each treatment group.

**Table 1.** Log Lung Fungal Burden between combination groups compared to fluconazole alone Negative binomial regression dispersion parameter 1.637 (theta), AIC 1817.7

| Experiment | Treatment Group | Fungal Burden Reduced | *Log Count Difference | p-value |
| --- | --- | --- | --- | --- |
| 1 | fluconazole:sertraline | yes | -1.5838 | $3.68 \times 10^{-06}$ |
| 2 | fluconazole:tamoxifen | yes | -2.4593 | $6.66 \times 10^{-13}$ |
| 3 | fluconazole:vanoxerine | No | 0.6742 | 0.0408 |

### Vanoxerine:fluconazole treatment is less effective than fluconazole alone

Mice were sacrificed on day 8. The average fungal burden for each group was: untreated infection 2.5 × 10^5^) (SD 1.5 × 10^5^), fluconazole alone 4.6 × 10^4^ (SD 3.4 × 10^4^), vanoxerine alone 4.6 × 10^5^ (SD 4.0 × 10^5^), and vanoxerine:fluconazole 9.3 × 10^4^ (SD 7.4 × 10^4^) total fungal colonies. Applying a negative binomial model, we discerned that combination therapy significantly increased the log fungal burden in the murine lung as compared to treatment with fluconazole alone (log increase 0.67, *P* <0.001, Table 1). Dissemination in the spleen was observed in the untreated infected group (16%), and in the fluconazole group (14%). Dissemination was not observed in the combination therapy group. Uninfected mice lost minimal weight during the treatment time (median, IQR; -0.3g, 1.56), whereas untreated infected mice lost weight (median, IQR; -4.1g, 2.3), fluconazole alone (median, IQR; -0.6g, 1.05), vanoxerine alone (median, IQR; -3.7g, 4.5), vanoxerine:fluconazole (median, IQR; -1.7g, 2.3).

### Sertraline:fluconazole treatment decreases fungal burden without impacting weight

Mice treated were euthanized on day 8. The average lung fungal burden for each group was: untreated infection 2.5 × 10^5^ (SD 1.5 × 10^5^), fluconazole alone 4.6 × 10^4^ (SD 3.4 × 10^4^), sertraline alone 2.7 × 10^5^ (SD 1.0 × 10^5^), and sertraline:fluconazole 9.8 × 10^3^ (SD 7.2 × 10^3^) total fungal colonies. Using a negative binomial model, we determined that combination therapy significantly reduced the log fungal burden in the lung when compared to fluconazole alone (log reduction -1.58, *P* < 0.001, Table 1). Dissemination in the spleen was observed in two groups: untreated infected (16%) and fluconazole alone (14%), but not in the combination therapy group. Healthy mice lost weight during the treatment time (median, IQR; 0-0.3g, 1.56), whereas untreated infected mice lost weight (median, IQR; -4.1g, 2.3). Fluconazole alone (median, IQR;-0.6g, 1.05) and sertraline:fluconazole groups gained weight (median, IQR; 0.5g, 1.1).

## DISCUSSION

Coccidioidomycosis remains a common infectious agent within endemic regions. Chronic or disseminated forms of the disease carry high associated morbidity, and in some cases may be fatal ^9,10^. Treatment durations range from weeks to months to life-long therapy; and shorter courses of therapy, or alternative treatment regimens in patients intolerant or refractory to current antifungals, are urgently needed.

The repurposing of currently available non-antifungal agents as adjunctive agents for coccidioidomycosis is a potential strategy to support available treatment options. Previously, we screened >1,000 FDA approved compounds for *in vitro* growth inhibition of *Coccidioides*. The agents evaluated in a murine model were selected based on availability of an oral formulation of the drug in question, the ability to achieve meaningful serum and tissue concentrations in a rodent model, and available toxicity data associated with systemic administration (tamoxifen, vanoxerine, sertraline).

Tamoxifen has anti-estrogenic activity and is used primarily in the treatment of early and advanced breast cancer. Off-target effects are common, and a number of such effects have been observed in mammalian cells ^11^. Additionally, tamoxifen possesses antifungal properties, as first noted against *Saccharomyces cerevisiae*^*12*^. Subsequent work has shown tamoxifen binding calmodulin and calmodulin-like proteins in *Cryptococcus* spp., which prevents the activation of calcineurin^11,13^, a key regulator of the yeast stress response; this results in cell lysis, decreased new bud formation, disruption of actin polymerization, and decreased germ tube formation^11^. The addition of tamoxifen to fluconazole treatment regimens resulted in fungicidal activity against *Cryptococcus* spp. *in vitro*^14^ and in a murine model of infection^13^, and human trials began shortly thereafter^15^. This open-label, randomized-control trial failed to demonstrate an increased rate of cryptococcal clearance from the cerebrospinal fluid of patients receiving this early combination therapy. However, tamoxifen remains a potentially attractive agent in the treatment of coccidioidomycosis because of the clear effects of estrogen and sex hormones on *Coccidioides* infections^16,17^. Tamoxifen concentrates in brain tissue (10-to 100-fold compared with plasma) and macrophage phagosomes, causing potential toxicity issues; and it is off-patent, potentially decreasing drug costs^15,18,19^.

The positive attributes prompted us to initiate *in vivo* experiments, despite potential drawbacks in clinical use. *In vivo*, tamoxifen:fluconazole treatments performed better than the current standard of care, fluconazole, (p<0.001) by log reduction of colony forming units (CFUs) in the lung. Additionally, there was no dissemination to the spleen in the combination treatment group; however, this was not significantly different from fluconazole or tamoxifen treatment alone. Interestingly, while tamoxifen treatment alone did not decrease fungal burden in the lung, we did see some evidence of protection in the spleen: only one tamoxifen-treated mouse displayed any fungal growth in the spleen, suggesting possible inhibition of dissemination. However, tamoxifen-treated mice were sacrificed early due to weight loss, clearly associated with treatment, exceeding 10% of starting body weight. In the control treatment groups, where mice received either tamoxifen alone or in combination with fluconazole, the mice also lost significant weight: an average of 11%. Tamoxifen interferes with glucose and lipid metabolism in mice,^20^ which is almost certainly the cause.

Vanoxerine is a dopamine re-uptake inhibitor and anti-arrhythmia medication previously evaluated in a number of clinical trials for these purposes^21–23^. Prior attempts to evaluate vanoxerine as an antimicrobial agent revealed antimycobacterial effects^24^. Disruption of the mycobacterial membrane, loss of membrane electrical potential, and the prevention of electrolyte efflux have been proposed as major mechanisms of vanoxerine’s antimycobacterial activity,^25^ and these same mechanisms may be responsible for its antifungal properties.

In our murine studies evaluating drug efficacy, vanoxerine:fluconazole combination therapy resulted in increased lung fungal burdens compared to fluconazole treatment alone. Although statistically significant, the p-value was just below the cutoff threshold (p-value = 0.0408). Dissemination was not seen in the combination group, suggesting that the addition of vanoxerine may have prevented extrapulmonary dissemination, yet increased the burden of pulmonary disease. The reasons for this are unclear.

Sertraline is a selective serotonin reuptake inhibitor (SSRI) used in the treatment of depression and anxiety disorders, among other conditions. Potent *in vitro* and *in vivo* fungicidal activity has been demonstrated against *Cryptococcus* spp., and the major mechanism of antifungal activity has been postulated as dose-dependent inhibition of eukaryotic translational initiation factor 3 (Tif3) by sertraline, with a resultant decrease in protein synthesis^26,27^. Based on these pre-clinical findings, a prospective, open-label, dose-finding pilot study was undertaken; and those receiving sertraline were found to have more rapid CSF cryptococcal clearance and a lower incidence of immune reconstitution inflammatory syndrome and relapse than was observed by similar cohorts in prior studies^28^. Seemingly contradicting this, a phase 3 study of HIV-infected patients with cryptococcal meningitis was disappointing, with no reduction in mortality or rate of fungal clearance from patient cerebrospinal fluid reported in those receiving adjunctive sertraline compared to placebo^29^. However, all patients in this trial received amphotericin B deoxycholate, and it is possible that its rapid fungicidal effects obviated any additive effects of sertraline. An alternative explanation might be that the time to steady-state of sertraline (7-14 days) was too long, for this brief study period, for a benefit to become apparent. In contrast, the treatment of severe forms of coccidioidomycosis require weeks to months of therapy, and in some cases are life-long. A delay in steady-state achievement of an adjunctive treatment agent is thus not detrimental and may still prove a useful addition.

In our murine model of infection, using a similar dosing strategy to that used in established models of cryptococcal infection, combination sertraline:fluconazole treatments performed better than the current standard of care, fluconazole (p < 0.001), as determined by measuring CFU in the lung. Additionally, there was no dissemination to the spleen in the combination treatment group; however, this was not significantly different from fluconazole treatment alone. Interestingly, sertraline treatment alone did not provide any protection: in both lung and spleen, similar burdens of infection were observed. All mice survived until the final day of the experiment. Weights were similar among all treatment groups, and all were higher than untreated mice, as expected.

In this study, we investigated the potential of three pharmaceutical agents, tamoxifen, vanoxerine and sertraline, to potentiate the fungistatic ability of fluconazole, the current standard of care in coccidioidomycosis. We observed that combination therapy with either tamoxifen:fluconazole or sertraline:fluconazole reduced fungal burden, in a murine model, compared to fluconazole alone. However, tamoxifen had a negative impact on the weight of the treated animals, necessitating early euthanasia, which in our opinion negates the drug’s potential antifungal efficacy. In contrast, sertraline:fluconazole-treated mice responded positively to treatment; this finding supports the feasibility of this therapeutic combination.

## Acknowledgements

The authors would like to thank Donald Chow for performing the small molecule screen for U.S. Patent 11,446,353 that led to this work.

---

**Supplemental Table 1.**
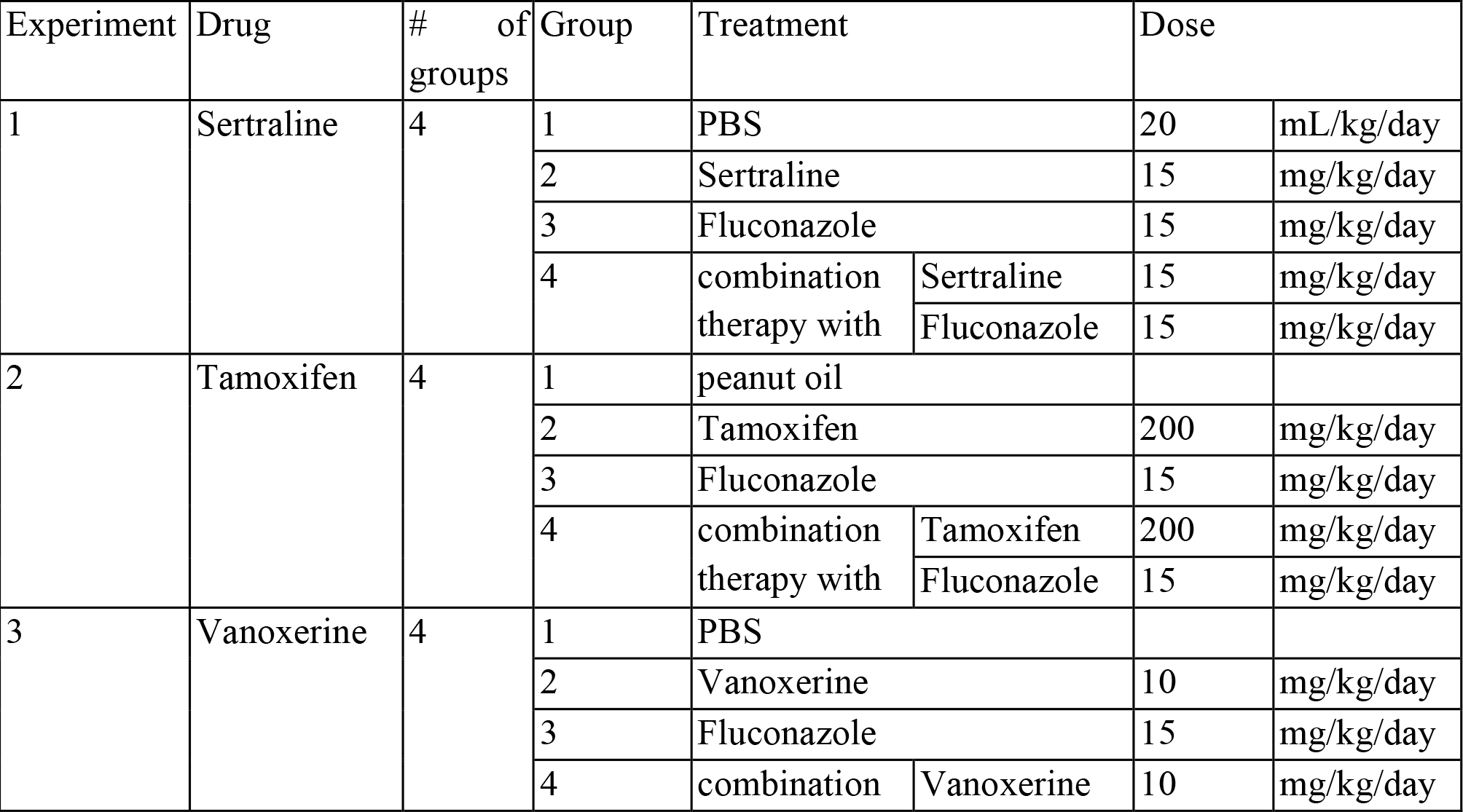
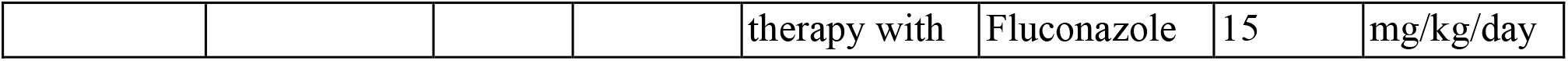
Breakdown of treatment groups.

